# COMPASS: An Open-Source, General-Purpose Software Toolkit for Computational Psychiatry

**DOI:** 10.1101/377556

**Authors:** Ali Yousefi, Angelique C. Paulk, Ishita Basu, Darin D. Dougherty, Emad N. Eskandar, Uri T. Eden, Alik S. Widge

## Abstract

Mathematical modeling of behavior during psychophysical tasks, referred to as “computational psychiatry”, could greatly improve our understanding of mental disorders. One barrier to broader adoption of computational methods is that they often require advanced programming skills. We developed the Computational Psychiatry Adaptive State-Space (COMPASS) toolbox, an open-source MATLAB-based software package. After specifying a few parameters in a small set of user-friendly functions, COMPASS allows the user to efficiently fit of a wide range of computational behavioral models. The model output can be analyzed as an experimental outcome or used as a regressor for neural data, and can be tested using goodness-of-fit methods. Here, we demonstrate that COMPASS can replicate two computational behavior analyses from different groups. COMPASS replicates and, in one case, slightly improves on the original modeling results. This flexible, general-purpose toolkit should accelerate the use of computational modeling in psychiatric neuroscience.

## Introduction

There is a growing need for advanced computational methods within psychiatric neuroscience (1, 2, 3). One particularly important aspect of that work is the development of quantitative, reproducible models that link patients’ symptoms to circuits, behaviors, and/or underlying psychological constructs (1, 2, 4). Such models have been used to quantify measurements of psychiatric disease processes relative to healthy subjects or to design experiments based on theoretical predictions (4, 5, 6, 7). Computational models are also expected to improve the reliability and utility of human neuroscience. For example, psychiatric neuroscience investigations usually classify subjects through categorical diagnoses and subjective rating scales. This, in turn, leads to poor signal-to-noise ratios (SNR) and difficulty in identifying reliable neural biomarkers of psychiatric illness (4, 8, 9). A more reliable approach may be to classify patients based on models of (ab)normal functioning. For instance, both patients and controls could perform the same standard psychophysical task, and patients’ degree of abnormality could be quantified based on the parameters of a model fit to their task behavior. Such models’ output(s) can become independent (regressor) variables in neuro-imaging or electrophysiologic analyses (10, 11, 12), potentially reducing inter-subject variability and improving SNR. Computational modeling also provides a framework for another major goal of psychiatric neuroscience: the identification of cross-diagnostic phenotypes and the circuits subserving those phenotypes (3, 9, 13, 14).

Modeling analyses of psychophysical task behavior often follow a common workflow. They cast the observed behavior as a function of an underlying theoretical construct, formalize that function in a system of parameterized equations, then identify parameters consistent with each experimental subject’s observed data. A challenge arises because each laboratory does this differently, often using custom-developed computer programs optimized for the modeling approach at hand. The resulting programs may not be well suited to analyzing slightly different datasets. Peer reviewers who are not modeling/programming experts are not readily able to assess whether the models were correctly implemented (15). Many researchers also do not have the mathematical/computational expertise to design such modeling systems *de novo.* Similar problems arose in the early days of neuro-imaging, and have been ameliorated at least in part by the development of freely available analysis packages that make it easier to apply best practices (15, 16, 17, 18, 19, 20, 22). Efforts exist to create similar analysis packages for behavior, e.g. hBayesDM (22) and KFAS (23). The available packages, however, do not work well with multiple behavior outputs (e.g., reaction times plus choices) and do not fully handle missing information in datasets (24).

Here, we present a general-purpose, open-source toolbox for fitting a wide variety of computational models to an equally wide variety of behavioral data. COMPASS is based on the state-space formalism, which assumes that behavior is influenced both by the parameters of individual task trials and by an underlying “cognitive state” that varies smoothly from trial to trial. This framework has successfully modeled behavior and neural activity in many contexts (4, 25, 26, 27), and fits the general concept that psychiatric symptoms arise from disruptions in basic underlying cognitive processes. Continuous (reaction times, physiologic measurements), binary (correct/incorrect, yes/no choices), and multinomial (learning of multiple stimulus-response contingencies) behavioral outputs can all be integrated into models, making the toolbox applicable to almost any laboratory task. To increase the applicability to “real world” data, COMPASS includes methods we recently developed to more optimally handle missing observations in these computational approaches (28).

We first provide a general overview of COMPASS, then demonstrate its application on two examples of associative learning behavior from published literature. In prior work, we showed how an early version of this toolbox could model reaction times in conflict tasks (4, 28). These examples illustrate the flexibility and generality of our approach. As a further illustration, the Supplementary Material shows the process of building a new model for a hypothetical decision task. Finally, a detailed user manual and code are available at https://github.com/Eden-Kramer-Lab/COMPASS.

### Overview of the state-space toolbox approach

Each step of the COMPASS analysis pipeline is implemented as a high-level function in MATLAB (MathWorks, Natick, MA; Figure 1). Because these steps are separated and scriptable, they can be configured to explore multiple models on a pilot dataset and determine which fits best before proceeding to hypothesis-driven analyses. The core assumption is that behavior is driven by a (potentially multivariate) cognitive state *X*, which varies over time according to its inherent dynamics and exogenous inputs:

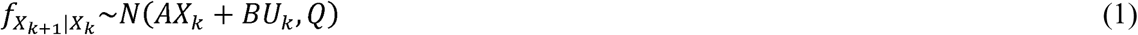

**Figure 1.**
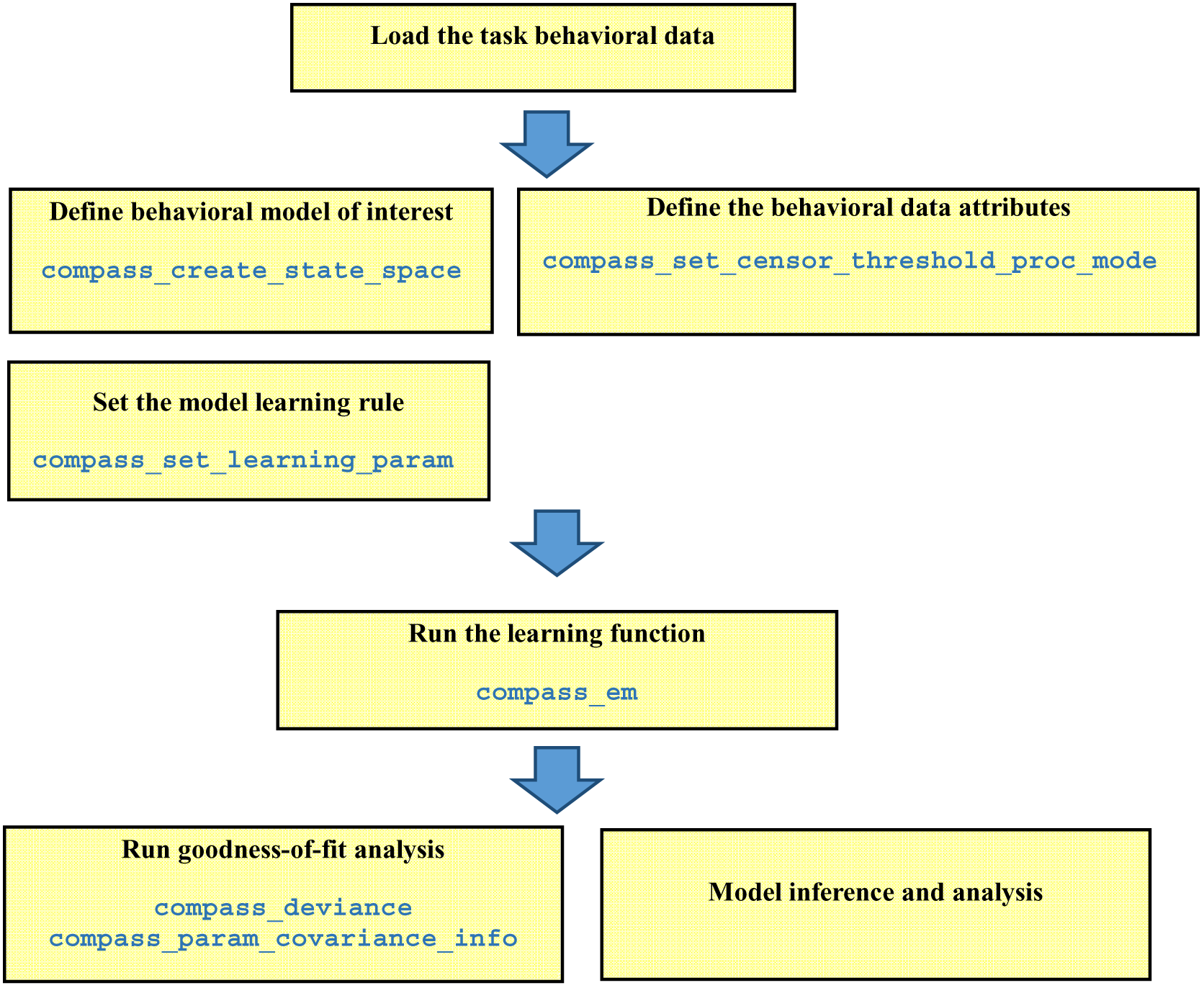
Pipeline of data analysis in COMPASS. The user manual at (https://github.com/Eden-Kramer-Lab/COMPASS/blob/master/compass_manual_march18_2018.pdf) describes each of these functions and how to use them.

That is, at time/trial *k*, the stated will evolve to its next value *X*_*k*+1_ based on innate transition dynamics *A* and its responsiveness *B* to an “input” *U_k_*. That evolution is driven by Gaussian noise with covariance equal to *Q*. The parameters for this model (*A, B, Q*) can be assigned by the investigator or inferred from the behavioral data. For instance, by assigning *Q* to have small diagonal elements and *A* to be a diagonal matrix with elements close to 1, we would obtain a state whose components are independent of each other and which will change very little trial-to-trial unless acted upon by *U_k_*. This might model a more “trait-like” process. Making *Q* elements larger would favor a process that changes substantially during an experiment, more “state-like”. The input *U_k_* may represent anything that would impact a subject’s performance, including features of the current trial, the outcomes of past trials (e.g., a running total of reward earned or punishments delivered), or the presence/absence of an external manipulation such as a drug or neural stimulation.

We cannot directly observe *X_k_*, but we observe its effects on the set of task-related behaviors *Y_k_*, which again may include non-conscious “behaviors” such as a physiologic response. This “observation process” follows a parametric distribution *g*:

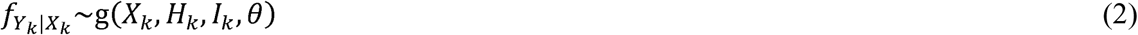

where *H_k_* represents the past history of the observations up to time *k*, and *I_k_*, like *U_k_*, may encode any other variables that can influence the behaviors *Y_k_* without having a persistent effect on the internal cognitive state *X_k_*. An example might be a conflict decision-making task, where trials with higher conflict/difficulty increase reaction times *Y*, but do not change the patient’s underlying decisional bias *X*(26). Many possible *Y* are not normally distributed. Binary choices usually follow a Bernoulli distribution, and both reaction times and physiologic outputs often follow a gamma or log-normal distribution (25, 26, 29). The COMPASS observation model thus allows us to model *Y* using a wide range of parametric distributions *g*, with parameters given by *θ*. We assume that *g* and *θ* do not vary trial-to-trial, and thus *θ* can be estimated directly from the data as with *A, B*, and *Q*. Further, *Y* may be multivariate, as in decision-making tasks where both the decision and the time needed to reach it may be experimentally relevant. In that situation, each element of the vector *Y_k_* may have its own distribution, described by a *g /θ* pair.

Choosing a model and assessing the goodness-of-fit of that model are critical components of any statistical analysis. Therefore, the toolbox provides a collection of different goodness-of-fit methods to help identify and refine models of the data. These include methods based on the covariance of the estimated parameters and the model deviance/likelihood. The online manual includes an example of those assessments for one of the task examples given below.

Another important analysis issue relates to the common experimental problem of missing data (30). Some data may be missing at random (MAR), such that the fact that a given datum is missing provides no new information. For example, response recording devices are subject to noise that may obscure some trials, such as muscle artifact in EEG or sensor failure for skin conductance. In such cases, trials with missing data are often removed prior to analysis. In other cases, features of the experiment influence the probability of data being missing. We identify these data as censored, or missing not completely at random. For example, in trial-based behavioral studies, subjects often fail to respond on individual trials within some designated time window, and this may be worse in patients taking psychotropic medications that slow processing. In such cases, it is inadvisable to simply remove trials with missing data, since these trials provide information about behavior. With censored reaction time (RT) data, we know that the subject’s RT was larger than a threshold, and this may affect the probability of a correct decision (28). When fitting a model with COMPASS, each observation in the matrix *Y* may be marked as observed, missing at random, or censored. COMPASS then incorporates this information in its state estimation and model identification processes, using algorithms described in (28).

### Example 1: Multivariate Associative Learning

Associative learning tasks are one of the most common models used to assess psychiatric deficits (2, 14, 31) and have been well-described using state-space models. In tasks where subjects must learn multiple associations simultaneously (32, 33), Prerau and colleagues described a method for inferring a single “learning state” (25). The learning state variable estimates how well the overall set of associations has been learned, optimally integrating performance over all available stimuli (25). The Prerau method also infers learning from both correct/incorrect choices and reaction times (RT), maximizing the information extracted from the available data (34, 35).

We analyzed a sample learning dataset from Williams & Eskandar (32) with both the original Prerau et al. code and an equivalent model set up in COMPASS (Figure 2). Here, we show an example from one behavior session, comprising 61 trials. On each trial, the subject (a rhesus macaque) attempted to learn/perform an arbitrary association between one of four stimulus pictures and four joystick directions. On 39 of the 61 trials, the subject indicated a choice within the required response window. For this example, the 22 non-response trials are excluded from the analysis. In the COMPASS user manual (see Supplementary Material) we show how to impute responses on non-observed trials. Thus, the behavioral signal processed in the toolbox includes the reaction time and decision over 39 trials (Figure 2A). The complete model is specified in only 10 lines of code, takes less than 20 seconds to run, and shows almost complete overlap with the output of the original authors’ custom code. We observe an inflection point around trial 15, where the subject’s learning state climbs rapidly as associations begin to be performed correctly (Figure 2B). The time to reach this or any other criterion point, or the slope of the learning curve around that point, could be used as a subject-level measurement of learning.

**Figure 2.**
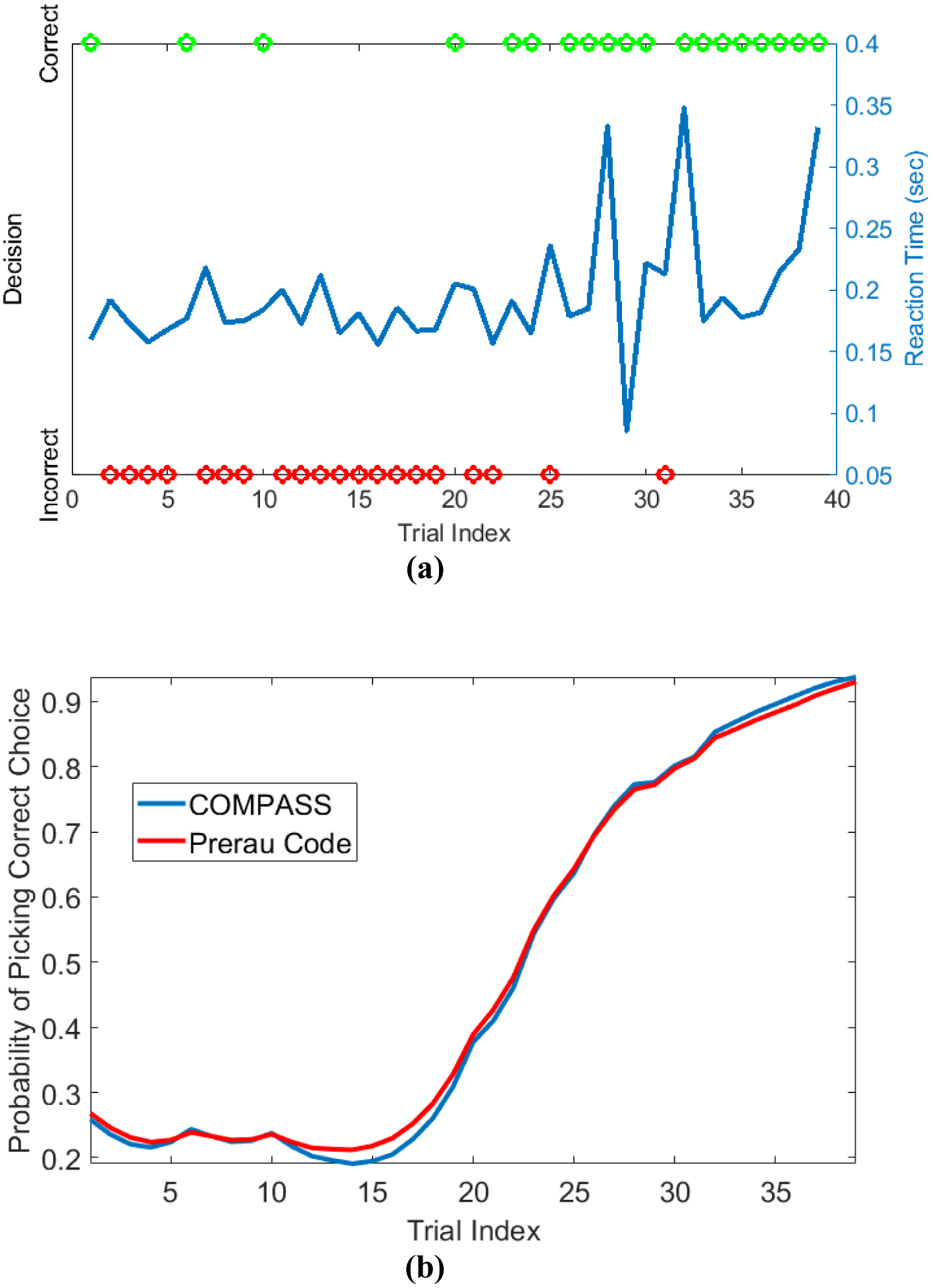
Sample learning behavior and learning state estimation using the method of Prerau et al. (26) and the COMPASS toolbox. (a) Correct (green) and incorrect (red) trial outcomes and reaction times (blue) over the course of a single task block. The behavior shows an abrupt change after 20 trials, with the subject beginning to match most stimuli correctly. The animal learns the task with an accuracy above 90% by the end of the trial block; the decision accuracy at the beginning of the block is less than 30%. (b) The estimated learning curve using the Prerau code (red) and the COMPASS toolbox (blue). The slight difference between the toolbox and Prerau et al. result relates to the different stopping criterion used in these methods. Prerau’s method sets a preset threshold on the likelihood growth and stops when the likelihood growth falls below the threshold. In COMPASS, *compass_em* stops after a user-specified number of iterations.

### Example 2: Negative Symptoms and Reward Motivation in Schizophrenia

Another major use of computational modeling is to tease out differential sensitivity to reward and loss, as in learning, gambling, and approach-avoidance tasks (6, 36). In one compelling example, Gold et al. (5) demonstrated this approach in schizophrenia, showing that patients with prominent negative symptoms were impaired at learning from gains, but not from losses. This was reflected in behavioral modeling, where high-negative-symptom patients specifically showed lower values of a model parameter reflecting how well they could compare gain-seeking to loss-avoiding options. The resulting analysis blended two common reinforcement-learning models, “actor-critic” and “Q-learning”, with the key parameter being how much each model drove or reflected behavior.

Appendix A shows how each term of the Gold et al. hybrid model can be captured in the state-space framework. In brief, the Gold et al. (5) task involves learning the correct option in four stimulus-action contingencies. We represent that learning progression by nine state variables; two state variables per contingency to represent the actor-critic learning process and one global state variable that represents the Q-learning process. The overall progress of learning for a given stimulus is given by a weighted combination of the state variables representing the actor-critic and Q-learning processes.

We analyzed a shared dataset, graciously provided by the authors, that is a subset of the original Gold et al. (5) dataset. On those data, using the Appendix A model, COMPASS replicated the original paper’s result. The data comprise 63 study subjects: 26 were healthy controls (HC) and the remaining 37 were clinically stable patients with schizophrenia or schizoaffective disorder. The latter were divided into high negative symptom (HNS, 19 patients) and low negative symptom (LNS, 18 patients) groups. Each subject performed 160 task trials, divided into 4 learning blocks of 40 trials. Behavioral outcomes included response accuracy (correct/incorrect), reaction time, money gained per trial, and trial type (Gain vs. Loss Avoidance).

Gold et al. (5) reported that HC and LNS subjects were more able to learn from gains than losses, while HNS subjects’ learning was more influenced by loss. This was reflected empirically in a greater accuracy on Gain than on Loss Avoidance trials (Figure 3a). It also was reflected in modeling and simulation of patients’ behavior at the end of task acquisition. When Gold et al. simulated data based on model parameters fit to each individual subject’s behavior, HC and LNS subjects were again more able to learn by obtaining gains rather than by avoiding losses (Figure 3b). We replicated this finding using COMPASS. Subjects’ behavior on our sample dataset showed the same pattern as in the original paper (Figure 3c). We then fit subject-level models to that behavior and plotted the individual subjects’ Gain vs. Loss Avoidance coefficients for accuracy prediction. This replicated the pattern of HC/LNS showing gain sensitivity and HNS showing primarily loss sensitivity (Figure 3d; Appendix A provides a detailed explanation of the Gold et al. computational model using COMPASS). In fact, the COMPASS modeling result is slightly more faithful to the empirical behavior pattern than the original Gold et al. simulation. The modeled performance of the HNS group (Figure 3d) is below the X-axis (as it is in the original empirical performance, Figure 3a), whereas the original simulations of Gold et al. produced mean Gain-Loss difference close to 0 (Figure 3b).

**Figure 3.**
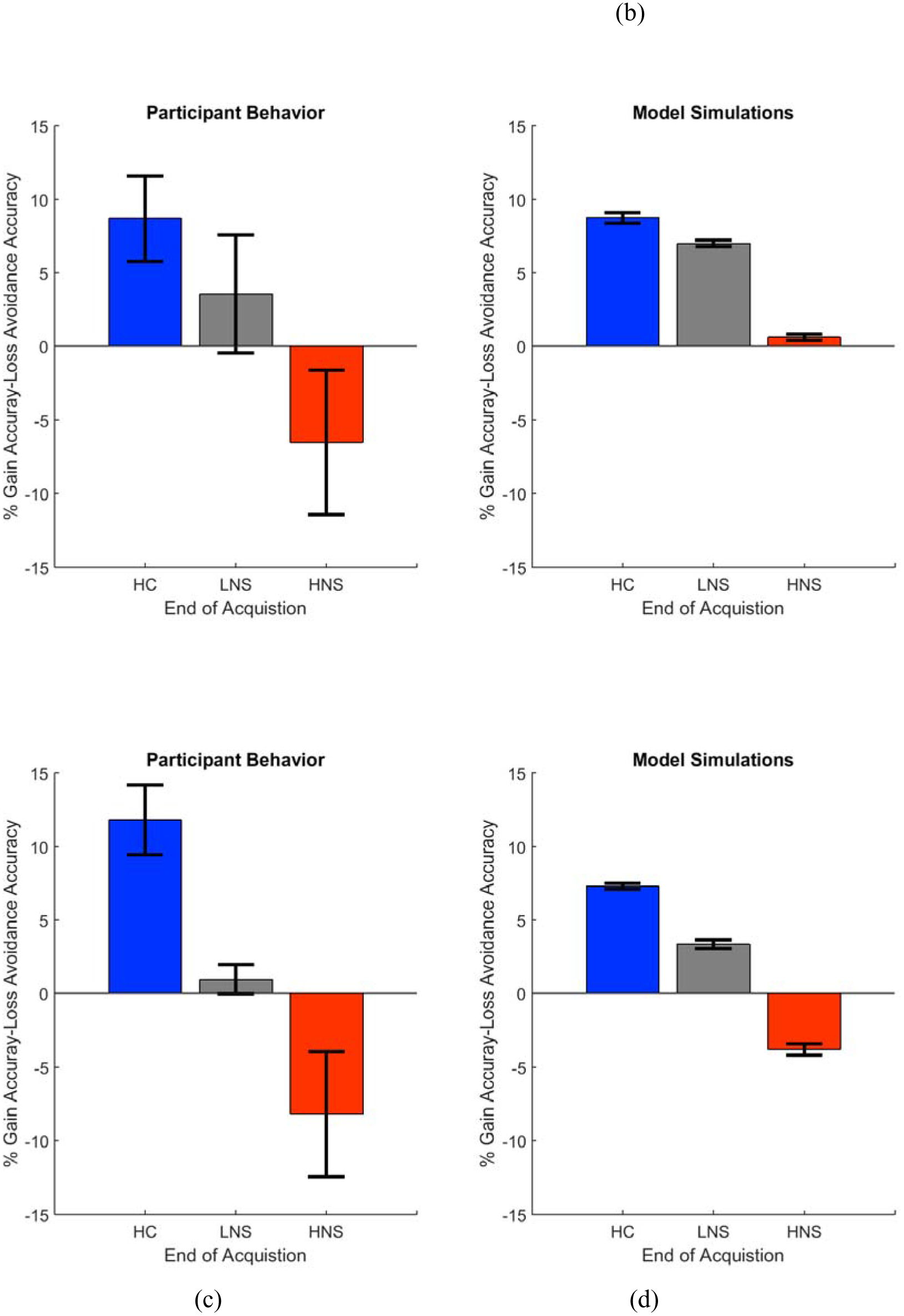
Observed (a, c) and simulated (c, d) end acquisition performance across patient and healthy control groups (HC, LNS, HNS). The HC group has a preference for learning from Gains (reward) relative to learning from avoided losses. This preference is reduced in the LNS group and inverted in the HNS group. The performance (% Gain accuracy - Loss Avoidance accuracy) is defined as the difference between the probability of picking the correct response on Gain trials and the probability of picking the correct response on the Loss Avoidance trials. (a) Observed performance and (b) simulated end of acquisition performance originally reported in Figure 4A-B of Gold et al. (2012). Simulated performance is generated from individual patients’ estimated model parameters, as described in the original paper. (c) Observed performance in the dataset shared by Gold et al. and (d) simulated result using an equivalent behavior analysis conducted in COMPASS. For (d), we run the equivalent hybrid model - described in Appendix A - per each patient. This gives, for each patient, an estimate (logistic regression coefficient) of the probability of making the correct choice on Gain vs. Loss Avoidance trials. The plot shows the average of these differences for each patient group. The pattern of HC > LNS > HNS on Gain-Loss Avoidance is replicated. Further, the COMPASS modeling simulation (d) matches the empirical behavior pattern (c) more closely than the original authors’ simulation.

## Discussion

We developed an open-source toolbox for state-space modeling and demonstrated its utility in analyzing dynamical behavioral signals relevant to computational psychiatry. In two examples from the learning literature, we showed that COMPASS replicates the results of prior modeling studies. In one of those examples, COMPASS estimated model parameters that more faithfully reproduced empirical results. The state-space modeling framework is a core tool in many research fields, especially engineering (37, 38). It is paving its way into neuroscience, psychiatry, and other medical research (39, 40, 41). COMPASS is the first attempt to provide unified and high-level functions that make a range of state-space models straightforward to implement and use for a wide variety of behavioral signals. Users can build a wide range of models to analyze behavioral data and compare their results in a principled way. A further explanation and guideline for building and comparing model forms can be found in the toolbox manual. The functions *compass_deviance* and *compass_param_covariance_info* enable model comparison. Further, COMPASS includes “on-line mode” functions that can continuously update state variables in real time as new data are acquired. The toolbox function *compass_filtering* addresses this, and examples of its use are given in the toolbox manual. In on-line operation, one can also call the learning algorithm periodically to re-update model parameters. Thus, COMPASS can also be used to construct experiments with adaptive behavior paradigms, where the stimuli presented to each subject are adjusted based on past performance. This approach could more efficiently sample regions of behavioral interest, e.g. the equipoise boundary of a choice task as modeled in (42, 43). It could also drive real-time application of a study intervention, e.g. brain stimulation or optogenetic (in)activation (4, 44).

The state-space modeling framework is not limited to normally distributed signals or discrete binary observations. COMPASS includes methods for highly skewed observation distributions (gamma and log-normal distributions) and for optimally imputing missing/censored data (26, 28). The distribution assumption is defined by arguments to the *compass_em* function, as are methods for censored data. These additions make COMPASS a powerful and versatile package for analysis of many different classes of dynamical signals. The main limitation of the state-space modeling framework is that prior to now, development and debugging of these models has been difficult. Development requires tedious work and extensive time, and involves statistical and programming skills that are not yet common in the field of cognitive neuroscience. We hope that by providing this toolbox, we can help other researchers delve into computational behavior analysis with a much lower barrier to entry.

## Acknowledgements

We thank Dr. Jim Gold, Dr. Michael Prerau, and their research teams for providing the data used in this manuscript. This research was funded by the Defense Advanced Research Projects Agency (DARPA) under Cooperative Agreement Number W911NF-14-2-0045 issued by ARO contracting office in support of DARPA’s SUBNETS Program. The views, opinions, and/or findings expressed are those of the authors and should not be interpreted as representing the official views or policies of the Department of Defense, the U.S. Government, or any other funder. The U.S. Government is authorized to reproduce and distribute reprints for Government purposes notwithstanding any copyright notation hereon.

## Appendix A Reformulating the Hybrid Learning Model Using the State-Space Modeling Framework

Gold et al’s. goal was to provide a quantitative fit to the pattern of data observed in patients and healthy controls. They investigated both a standard Actor-Critic architecture and a Q-learning architecture. They argue that neither taken alone could account qualitatively for both healthy control and patient data. They thus investigated a mixture model of Actor-Critic and Q-learning, which led to better qualitative and quantitative fits for all groups and explained key features of the data. The actor-critic model uses reward prediction errors to modify the probability of selecting an action, while the Q-learning model predicts sensitivity to actual outcome values. That is, Q-learning is more focused on maximizing total value, and therefore predicts that subjects will choose a high chance of gain over a high chance of avoiding a loss. Actor-critic would value those two choices equally, because they have similar chances for prediction error. Given that different patient groups showed a mixture of these strategies in responding to the task, the hybrid model should be able to better account for observed results in patients and healthy subjects.

Here, we show how the model of Gold et al. (5) can be implemented using the state-space modeling framework and thus the COMPASS toolbox. Table A.1 shows a line-by-line comparison between two models. Here, we focus on the Hybrid model proposed in the paper. The pure actor-critic or Q-learning processes are special cases of this model with parameter *c* (see below) set to 0 or 1, respectively.

**Table A.1.**
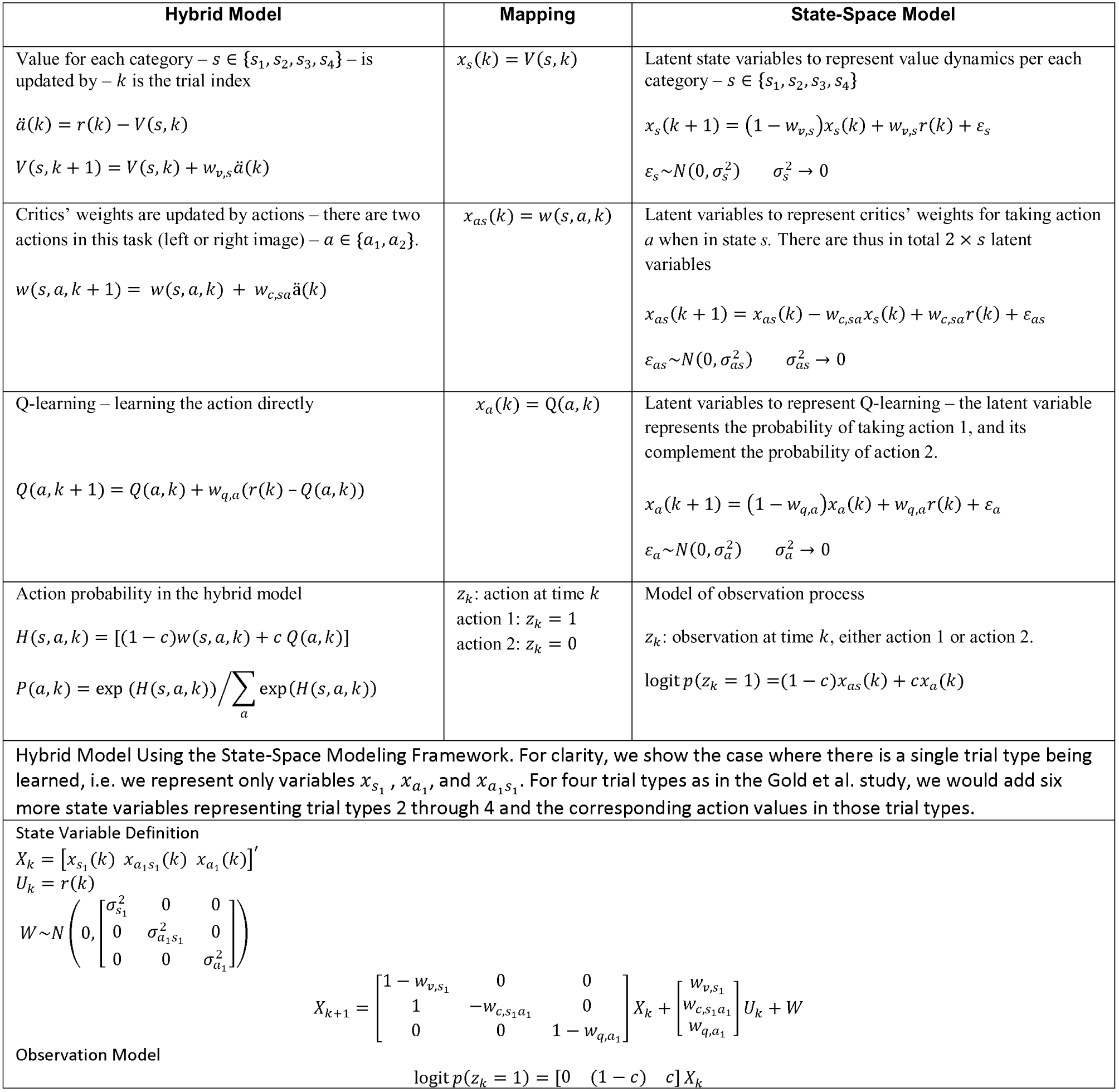
Hybrid model of actor-critic and Q-learning and its equivalent state-space model

The model is cast in terms of the trial type – *s* – and the possible actions within that trial type – *a. a* can be seen as picking one of two actions, where a correct action on the gain trials corresponds to picking the item associated with winning the reward. The correct action on the loss-avoidance trials corresponds to picking the item that is not associated with a penalty. Note that we only need to represent one of these two actions per category with a state variable; this is because not taking an action implies that the other action is being taken. We assign two state variables to each task’s category to represent the value of being in a state and of taking action 1 during that state. That is, we need *x*_*s*_1__ and *x*_*a*_1_*s*_1__ to present the probability of the stimulus being category 1 and the probability of taking the action *a* when in category 1. We assign one state variable to represent the global value of taking action 1 across all possible states/trial types – *x*_*a*_1__ (*k*). Now, we can define the state-transition process for the task by (note that action *a*_1_ will be common across states):

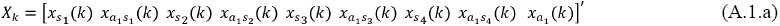

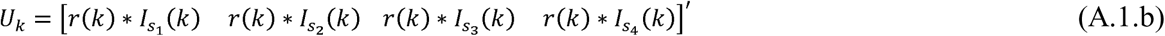

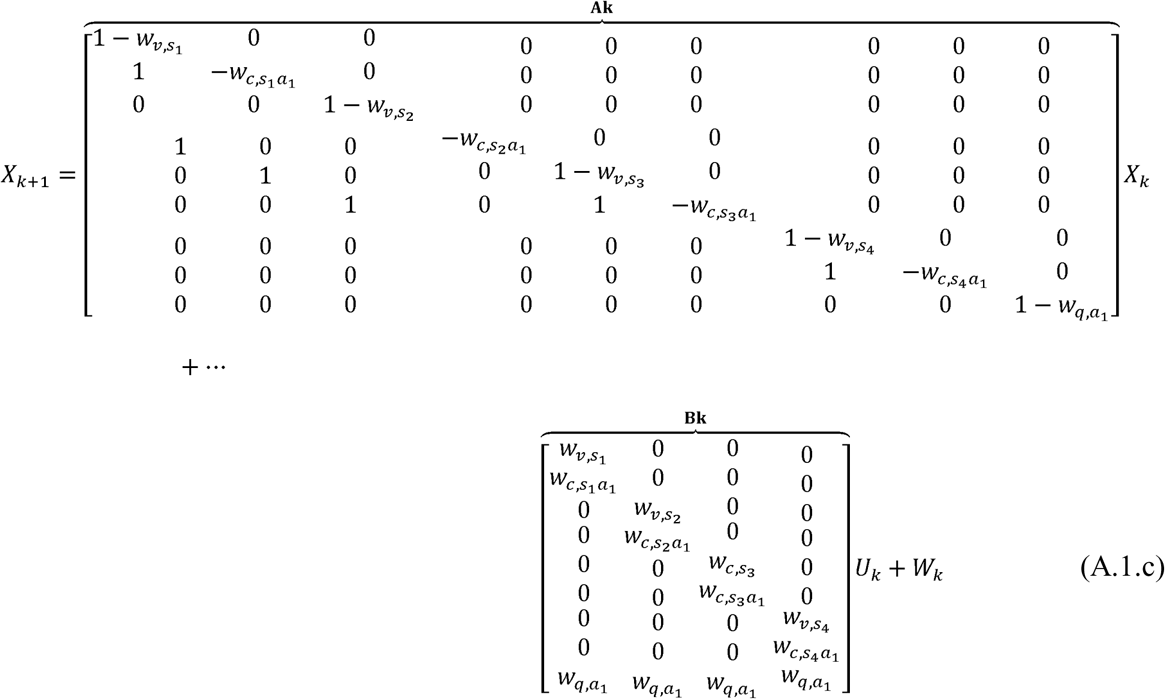

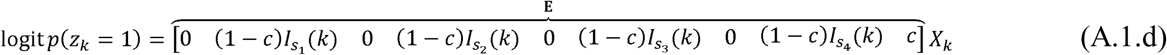

where, *_s_i__* is an indicator function which is 1 when the current trial type corresponds to stimulus category *i*. In equation (A.1.c), the probability of taking each action is defined as a function of the learning, captured in the vector *X_k_*. *X_k_* represents both actor-critic and Q-learning values - the first 8 entries describe actor-critic learning for each trial type, while the 9^th^ element represents the global Q-learning process that focuses solely on the action regardless of the trial type. As Gold et al. did, we represent the relative weight of these two learning processes (actor-critic and Q-learning) through a mixing parameter *c.* Using the Logit function defined in (A.1.d), the probability of taking action *a*_1_ is defined by *c x*_*a*_1__(*k*) in the absence of the actor-critic model. The probability of taking action *a*_1_ when presented with trial type *s*_1_, in the actor-critic model, is defined by (1 – *c*)*x*_*a*_1_*s*_1__(*k*). The overall probability of taking action *a*_1_ when presented with *s*_1_ is thus defined by *c x*_*a*_1__(*k*) + (1 – *c*)*x*_*a*_1_*s*_1__(*k*). We can similarly define the probability of both actions for the other 3 trial types. Given the model definition, an increase in *x*_*a*_1__(*k*) implies a global preference for *a*_1_ in all trial types, while an increase in *x*_*a*_1_*s*_1__(*k*) implies a specific preference for *a*_1_ when presented with *s*_1_.

The model of Table A.1 translates directly into MATLAB code, as shown in Figure A1. This close mapping between model specification and computer code makes COMPASS easier to use and interpret.

**Figure A.1.**
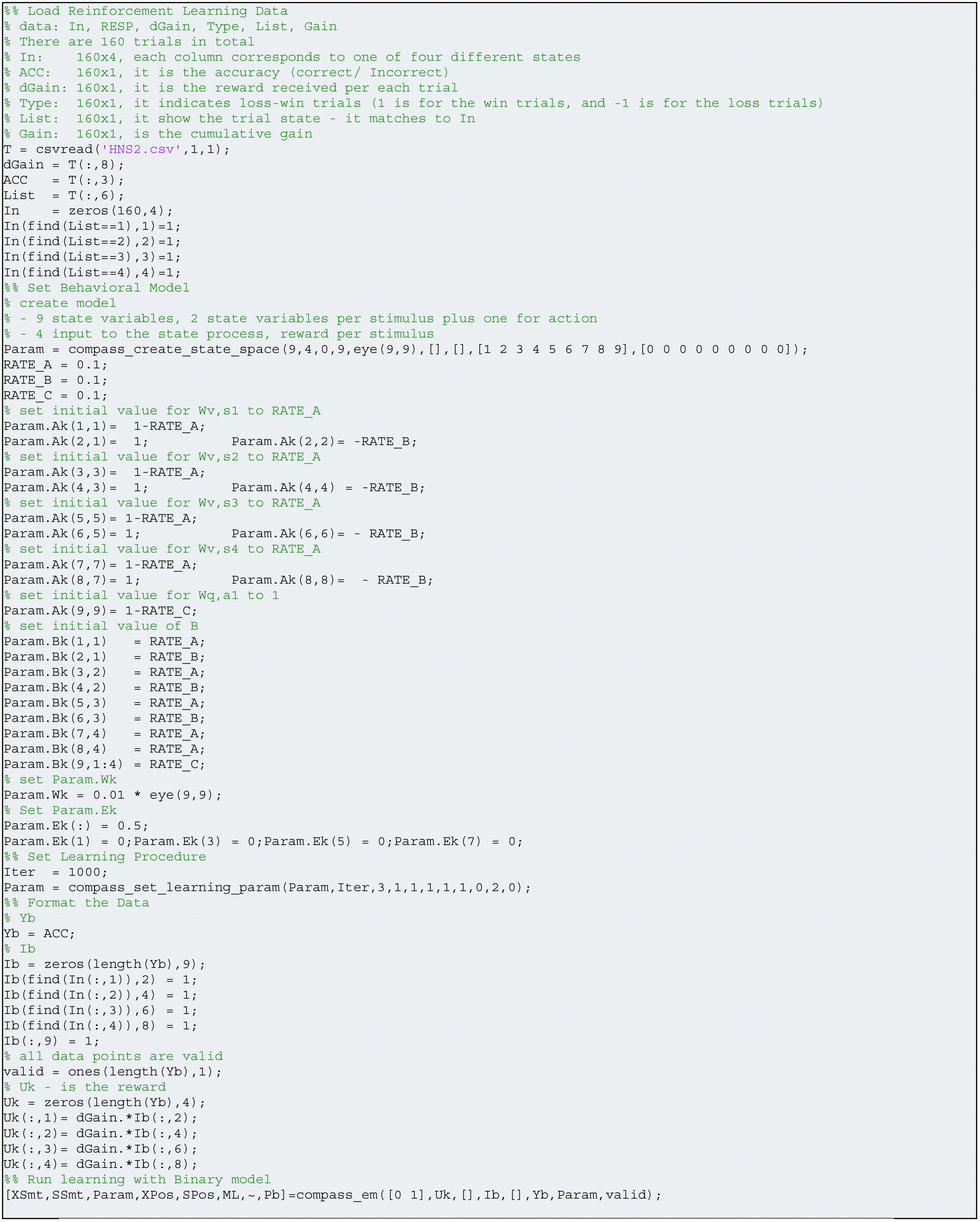
Analysis script for the hybrid learning task

In the script, RATE_A and RATE_B define initial values for *w_v,s_i__* and *w*_*c,s*_*i*_*a*_1__ *i* = 1 ··· 4. RATE_C defines an initial value for *w*_*q*, *a*_1__. The initial values for RATE_A, RATE_B, and RATE_C are set to 0.1. This setting defines initial values for elements of *A* and *B* matrices, which defines how new reward information are combined with the previous estimation of state variables. The optimal values for *w_v,s_i__*, *w*_*c,s*_*i*_*a*_1__ and *w*_*q,a*_1__ are smaller than 1 and preferably close to zero, if the values for the critics’ weights and values dynamics (**Table A.1**) reasonably replicate the task participant behavior. As a result, we initialize these parameters with a small number and then allow the E-M algorithm to drive them to their optimal values. The *c* parameter, defined in (A.1.d), is set to 0.5 in the script and then updated by being learned from each subject’s data. We define a covariance matrix for *W_k_* using *Param. Wk*. This is set to be a diagonal matrix with a small variance, because the terms are independent of each other and there is an overall assumption that we can predict behavior accurately if these variables are known. Values of *Param.Ek* are set to either 0 or 0.5; it is set 0 for index 1, 3, 5, and 7 and it is set to 0.5 for index 2, 4, 6, 8 and 9. Note that *Param.Ek* is fixed and it is not adjusted in *compass_em.* The settings suggest that a mixture model (actor-critic and learning) is being trained in the model. If we set *Param.Ek*(9) to 0, we then have a model solely based on actor-critic. If we set all elements of *Param.Ek* zero except *Param.Ek*(9), we then build a model based on Q-learning. The input vectors *Ib* and *Uk* represent the stimulus information and obtained reward on each trial for the subject being analyzed.

The behavioral signal analyzed in Gold et al. is solely the action – or decision –taken on each trial. However, there is also a reaction time signal, which might carry extra information about the learning evolution or attribute, as in the Prerau et al. papers discussed in section 4. For instance, we might expect to see a shorter reaction time as the correct actions are learned and the subject gains confidence in his/her decisions. It would be straightforward to add reaction time to the model as a function of the learning variables, *X_k_*, or to add the previous decision outcome as a history term. We can define how the reaction time is linked to the model state variables or input in the function *compass_create_state_space*, and pass both continuous and discrete inputs to the *compass_em* routine. We can also define how parameters of the reaction time model will be trained using *compass_set_learning_param*. We even can define the censoring criteria if some data points are censored due to long response times, *compass_set_censor_threshold_proc_mode*.

